# Black Soldier Fly Bioconversion to Cultivated Meat Media Components Using Blue Catfish Gut Microbiome

**DOI:** 10.1101/2024.01.20.576372

**Authors:** Alfonso Timoneda, Arian Amirvaresi, Reza Ovissipour

## Abstract

Developing low-cost media is one of the major challenges in the cellular agriculture domain. Thus, this study aimed to develop low-cost media for cell-cultivated seafood using gut-microbial community-assisted fermentation. Black soldier fly larvae (*Hermetia illucens*) were used as the substrate and exposed to gut microbial communities isolated from Blue catfish (*Ictalurus furcatus*). In the first step, BSFL slurry was subjected to enzymatic digestion, using pepsin and trypsin to mimic animal digestive processes. The results showed a 2.8% degree of hydrolysis after digestion with pepsin and an additional 5.9% after digestion with trypsin. In the second step, two fermentation approaches were tested, including the direct addition of gut homogenate to the hydrolysates (fermentation A) and the establishment of microbial cultures from the gut homogenate before fermentation (fermentation B). Both fermentations resulted in similar protein content and degree of hydrolysis. Fermentation led to a decrease in species richness, with the loss of important chitinase and protease-producing genera such as *Pseudomonas* and Clostridiaceae. However, there was an increase in *Paraclostridium* and members of the Enterobacteriaceae. In addition, the effect of fermented hydrolysates from BSFL on the proliferation of zebrafish embryo fibroblasts was tested in comparison to fetal bovine serum (FBS) in in vitro cell cultivation. Lower concentrations of FBS resulted in decreased cell density and altered cell morphology. The supplementation of hydrolysate B at high peptide concentrations had cytotoxic effects on the cells, while at lower peptide concentrations, it improved cell proliferation only in cultures with 2.5% FBS.

## Introduction

Cellular agriculture is a burgeoning and revolutionary field that has the capability to tackle conventional farming challenges. By leveraging breakthroughs in cellular and biotechnological advancements, it aims to generate animal-derived products without relying on traditional farming methods (Batish et al. 2022; Nikkhah et al. 2023; Stout et al. 2023). The successful large-scale commercialization of cultivated meat continues to face a significant technical challenge in the formulation of cell culture media (O’Neill et al. 2023; Stout et al. 2023). The primary challenge in this domain is the development of serum-free media that are both cost-effective and efficient. Researchers and industry players are actively addressing this problem by exploring the utilization of plant-based (Stout et al. 2023; Yamanaka et al. 2023), microbial and insect-based proteins (Batish et al. 2022). To mitigate the cost of media, a promising approach involves leveraging low-cost resources, including agricultural wastes, or exploring emerging protein sources such as insects that have been reared with agricultural waste. Recently, we have successfully developed low-serum and serum-free media utilizing protein isolate and enzymatic-assisted protein hydrolysates from black soldier fly larvae (Batish et al. 2022). While protein extraction through chemical and enzymatic digestion methods yields proteins suitable for replacing serum in cell culture media, these processes entail a considerable use of chemicals, enzymes, thermal processing, and energy. Moreover, these processing methods are selective, extracting only soluble proteins and peptides during chemical or enzymatic extractions from the feedstock. However, the feedstock comprises a significant amount of lipids, carbohydrates, fibers, and minerals, which could potentially be converted into valuable metabolites for cell culture.

*In vivo*, foods undergo exposure to various enzymes and microbial fermentation within the gastrointestinal system. This process leads to the generation of short-chain fatty acids, peptides, exopolysaccharides, cell surface proteins, and beneficial enzymes that contribute to the overall well-being of the host (Youn et al. 2022). Additionally, the fermentation process and microbial conversion of feedstock into value-added products is a green and clean approach (Gadkari et al. 2021). Our team efforts have been mainly focused on protein hydrolysates and protein isolate applications to replace or reduce FBS in cultivated meat media (Batish et al. 2022). Based on our previous studies, we concluded that black soldier flies larvae protein hydrolysates and protein isolated at low concentrations support cell proliferation.

In this study, we were inspired by nature and utilized the gut microbial communities of Blue catfish (*I. furcatus*) for black soldier fly larvae bioconversion. The resulting postbiotics were then employed as a substitute for fetal bovine serum (FBS) in Zebrafish fibroblast cell culture media. This innovative approach leverages the inherent digestive processes of catfish microbiota to convert black soldier fly larvae into valuable postbiotics. The selection of Blue catfish was deliberate, given its status as one of the most abundant invasive fish species in the U.S., primarily feeding on insects.

## Material and Methods

### Enzymatic hydrolysis of black soldier fly larvae

Black soldier fly larvae (BSFL) were acquired as a dried sample (Sun Grubs, USA). BSFL were blended to a flour and mixed with deionized water in a 1:3 ratio (w/v). The resulting slurry was mixed for 2 h at 37**°**C in a precision digital circulating water bath (Thermo Fisher Scientific, Waltham, MA, USA) with a rotator blade at 150 rpm. To best mimic the digestion procedure in animal digestive tracts, the BSFL slurry was sequentially digested first with pepsin from porcine gastric mucosa (Sigma-Aldrich, Saint Louis, MO, USA) at pH 1.2 for 1 h, and later with trypsin from porcine pancreas (Sigma-Aldrich, Saint Louis, MO, USA) at pH 8 for 1h, both at 37**°**C in a water bath with a rotator blade at 150 rpm. Both enzymes were added at a 2% ratio of enzyme to total protein. After that, enzymes were heat-inactivated by bringing the slurry to 100**°**C for 10 min. Samples were taken before, between, and after enzymatic digestions and stored at -20°C for later assessment of the degree of hydrolysis. The initial water content of BSFL was assessed by the weight difference of 1 g of BSFL before and after 52 h of a drying process at 50**°**C (N=2).

### Blue catfish gut dissection

The blue catfish (*Ictalurus furcatus*) used for gut dissection was acquired early in the morning at Amory Seafood Co. (Hampton, VA, USA) and was originally fished in the Pasquotank River (NC, USA) less than 24 h prior, in April of 2023. The fish weighed 539 g and was kept on ice from capture until dissection. Dissection was performed in sterile conditions in a Thermo Scientific 1300 Series Class II, Type A2 biological safety cabinet (Thermo Fisher Scientific, Waltham, MA, USA). The fish ventral area was disinfected three times by rubbing with 70% ethanol before incision. The whole gut was extracted and cut from the start of the midgut to the end of the hindgut. The gut contents were squeezed out and discarded, and the intestinal tube was washed with deionized water. The extracted gut weighed 6.20 g and was transferred to 22.5 ml of sterile Tryptic Soy Broth (TSB). Gut was homogenized with a Fisherbrand 850 tissue homogenizer (Thermo Fisher Scientific, Waltham, MA, USA) for 4 min at 12000 rpm.

### Anaerobic fermentation of BSFL hydrolysates

Two approaches for the fermentation of BSFL hydrolysates with fish gut microbiome were assayed. For fermentation A, 10 mL of gut homogenized was directly added onto the BSFL hydrolysate and incubated in anaerobic conditions at 29**°**C and 150 rpm for 24 h. Anaerobic conditions were achieved by using GasPak™ EZ Gas Generating Pouch Systems containing anaerobic blue/white indicators (Becton Dickinson, Crystal Lake, NJ, USA) and double bagging each culture with plastic zip bags. For fermentation B, microbial cultures were established from the homogenized gut before fermentation. Three different media conditions were assayed: TSB at a 1:10 ratio (v/v), TSB with 0.5% of yeast extract at a 1:10 ratio (v/v), and TSB with 0.5% of yeast extract at a 1:5 ratio (v/v). Cultures were grown in anaerobic conditions at 29**°**C and 150 rpm for 24 h, and bacterial growth was assessed by measuring cultures’ absorbance at 600 nm using a Biotek Epoch microplate spectrophotometer (Agilent, Agilent, Santa Clara, CA, USA). Cultures in TSB with 0.5% of yeast extract at a 1:10 ratio (v/v) were concentrated 5-fold by a 5 min centrifugation at 500 *g* and resuspension of bacterial pellets in 10 mL of supernatant. Fermentation was established by adding the bacterial concentrate onto the BSFL hydrolysate and followed by incubation in anaerobic conditions at 29**°**C and 150 rpm for 24 h. Harvest of the fermentation products and non-fermented controls was performed by centrifugation of 5 min at 5000 *g*, and sequential filtrations of the liquid phase through 100 μm, 70 μm and 40 μm cell strainers (Thermo Fisher Scientific, Waltham, MA, USA) and a final sterilization step with a Millipore 0.22 μm stericup vacuum filter system (Sigma-Aldrich, Saint Louis, MO, USA). Samples for analysis of bacterial communities were taken before and after fermentations, and after bacterial growth in TSB with 0.5% of yeast extract at 1:10 ratio (v/v). Fermentation products were stored at -80**°**C until use.

### Taxonomical evaluation of bacterial communities

Identification and quantification of bacterial genus in samples taken throughout the fermentation process was performed by CD Genomics (Shirley, NY, USA). The microbiome total DNA was extracted using the QIAamp DNA Microbiome Kit (Qiagen, Hilden, Germany) according to the manufacturer’s protocol. The quality and concentration of the extracted DNA were measured using a ND-1000 NanoDrop spectrophotometer (NanoDrop Technologies, Wilmington, DE, United States). Microbial diversity was assessed by sequencing of the 16S ribosomal RNA amplicon on a paired-end Illumina MiSeq platform. Paired-end reads were merged using FLASH (V1.2.11). Quality filtering of raw tags was performed according to the Fastp quality control process. Operational Taxonomic Units (OTU) were clustered and taxonomically assigned with QIIME 2, using a pre-trained Naive Bayes classifier. Heatmap analyses were carried out and plotted using R language tools and Ward hierarchical clustering. Four different metrics were calculated to assess the alpha diversity: Chao1, Ace, Shannon and Simpson.

### Protein and peptide quantification

Protein quantification was performed using the Pierce Detergent Compatible Bradford Assay kit (Thermo Fisher Scientific, Waltham, MA, USA) following manufacturer’s specifications. In brief, 10 μl of sample was mixed with 300 μl of Bradford reagent in 96 well plates, incubated 10 min at room temperature, and measured at 595 nm with a Biotek Epoch microplate spectrophotometer (Agilent, Agilent, Santa Clara, CA, USA). A standard curve with Bovine Serum Albumin (BSA) was performed in a range between 0 and 750 μg/mL. Every measurement was performed in triplicate.

The degree of protein hydrolysis was measured using the *o*-phthalaldehyde (OPA) assay, based on the reaction of OPA and 2-mercaptoethanol with amino groups released during proteolysis of a protein substrate (Church et al. 1985). Fresh OPA solution was prepared as follows: Solution A, consisting of 7.62 g of sodium tetrahydroborate and 200 mg of sodium dodecyl sulfate (SDS) in 150 mL of DI-water, was first mixed with solution B, consisting of 160 mg of o-phthaldialdehyde in 4 ml of methanol 100%. The resulting solution was then mixed with solution C, consisting of 400 μl of 2-mercaptoethanol in 50 mL of DI-water. 270 μl of OPA solution was mixed with 36 μl of sample in 96 well plates and incubated at room temperature and darkness for 2 min. Absorbance of each sample was measured at 340 nm in a Biotek Epoch microplate spectrophotometer (Agilent, Agilent, Santa Clara, CA, USA). Every measurement was performed in triplicate. For peptide quantification, a standard curve was performed with serine from 0.0625 to 0.5 mg/mL. The degree of hydrolysis was calculated using equation 1. Total hydrolysis of the original sample was obtained by acid hydrolysis with 6 N HCl at 110**°**C for 24 h.

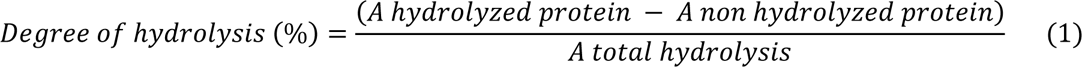

### Cell culture and maintenance

The cell line used was the zebrafish (*Danio rerio*) embryo fibroblast ZEM2S cell line and was originally obtained from American Type Culture Collection (ATCC) (Manassas, VA, USA)(Batish et al. 2022). ZEM2S cells were grown in T75 flasks with 10 mL of cell maintenance media composed of Leibovitz L-15 media (L-15) with 10% fetal bovine serum (FBS) and 1% antibacterial-antimycotic. Cells were incubated in a 1555 Shel Lab incubator (VWR, Radnor, PA, USA) at 27**°**C. Subculturing was performed when cells reached a confluency of 80-85% by removing media, rinsing cells with phosphate-buffered saline (PBS) and detaching cells from the flask surface with trypsin for 3-4 min. Trypsin was then neutralized with cell maintenance media and cells were recovered by centrifugation for 8 min at 500 *g*. Cell pellets were resuspended in 2 mL of cell maintenance media and quantified using a Countess automated cell counter (Thermo Fisher Scientific, Waltham, MA, USA). Seeding density for cell maintenance was approximately 9,300 cells/cm2. Cell growth and morphology were routinely assessed using an Olympus CKX53 phase-contrast microscope equipped with an Olympus DP27 image camera and the Olympus cellSense Entry imaging software (Olympus, Tokyo, Japan).

### Cell proliferation assessments

For ZEM2S cell proliferation experiments cells were cultured in 24-well plates at an approximate seeding density of 10,000 cells/cm^2^ with cell maintenance media (L-15 with 10% FBS and 1% antimicrobial-antimycotic) for 24 h to allow for cell adhesion. Cells were grown in a 1555 Shel Lab incubator (VWR, Radnor, PA, USA) at 27**°**C. After 24 h, cell media was replaced by the assayed media treatments, which combined different FBS concentrations (10%, 2.5%, 1%, 0%) and peptide concentrations (0.01 mg/mL, 0.001 mg/mL, 0 mg/mL) of fermentation products A and B, and no-fermentation controls (C-A and C-B). Control A and B are the product of a BSFL hydrolysate that did not undergo bacterial fermentation with blue catfish gut microbes, without and with 10% of TSB with 0.5% of yeast extract, respectively. In all cases, pH of fermentation products and hydrolysate controls were re-adjusted to 7.8 before supplementation to cell culture media. Cells were grown in each treatment for 3 days with four replicates per treatment.

Cell number and morphology were assessed at times 0 h, 24 h, 48 h and 72 h, from the moment of media replacement. Cell quantification was performed using the CKX-CCSW confluency checker software (Olympus Life Science, Waltham, MA, USA) by taking three images per well. Cell proliferation was assessed for each well as the fold-change increase in cell count calculated over the first time point (0 h). Values for each biological replicate were averaged and error was calculated as the standard deviation.

### Statistical analysis

Statistical differences in each experiment were assessed via one-way ANOVA analysis of variance and Tukey’s Honest Significant Difference test.

## Results and Discussion

### Enzymatic digestion of BSFL

BSFL were sequentially subjected to enzymatic digestion and bacterial fermentation in an attempt to mimic conditions for protein breakdown naturally presented in the fish digestive tract. The initial water content of BSFL was 3%, thus BSFL was blended and mixed with DI-water. The resulting BSFL slurry was subjected to a 1 h digestion with pepsin at pH 1.2, followed by another 1 h digestion with trypsin at pH 8. The degree of protein hydrolysis was assessed throughout the process by using the *o*-phthaldialdehyde (OPA) spectrophotometric method, which accurately detects primary amines present in amino acids and peptides released through proteolysis (Church et al. 1985). Digestion with pepsin resulted in a 2.8% degree of protein hydrolysis, whereas digestion with trypsin added up to a further 5.9% (Table 1). These results are in the range of previously reported BSFL enzymatic digestions showing a 2-3% degree of hydrolysis after 1 h with alcalase (Ravi et al. 2020), and a 6%, 17%, 25% and 25% after 16 h with protease from *Bacillus licheniformis*, pepsin, papain and pancreatin respectively (Caligiani et al. 2018). However, they are lower than the 20% reported after 1 h digestion of BSFL with alcalase from *B. licheniformis* (Batish et al. 2022). This could be attributed to different enzyme efficiencies or the protein separation process samples underwent after enzymatic digestion in their case (Batish et al. 2022).

**Table 1.** Degree of hydrolysis of BSFL during the enzymatic digestion and fermentation process^1^.

| Stage of hydrolysis | Peptides<br>(mg/ml) | Degree of hydrolysis<br>(%) |
| --- | --- | --- |
| BSFL slurry | 9.3 | 0 |
| BSFL slurry + pepsin | 13.3 | 2.8 |
| BSFL slurry + pepsin+ trypsin | 17.8 | 5.9 |
| BSFL slurry + pepsin+ trypsin+ fermentation A | 34.3 | 15.3 |
| BSFL slurry + pepsin+ trypsin+ fermentation B | 35.5 | 16.1 |
| BSFL slurry + HCL 6N (total hydrolysis) | 205.7 | 95.7 |

<sup>1</sup> Subsequent digestions with pepsin and trypsin were both performed at 37°C for 1 h and at pH 1.2 and pH 8, respectively. Total hydrolysis of the original sample was obtained by acid hydrolysis with 6 N HCl at 110°C for 24 h

### Fermentation of BSFL with blue catfish gut microbiota

After enzymatic digestion, the resulting BSFL slurry underwent bacterial fermentation with gut microbes isolated from blue catfish (*I. furcatus*) (Figure 1). In fermentation A, a portion of the blended gut was directly added to the BSFL slurry and incubated for 24 h in anaerobic conditions. This resulted in 34.3 mg/mL protein, with 15.3% degree of hydrolysis (Table 1). While in fermentation B, in which we attempted to establish microbial cultures from samples of blended gut 24 h before fermentation by using various liquid media conditions, protein content was 35.5 mg/mL, with a degree of hydrolysis of 16.1% (Table 1). The results indicated that there were no significant differences between Fermentation A and B. In fermentation B, bacterial cultures were grown for 24 h in anaerobic conditions, and bacterial growth was assessed by the increase in OD600. Cultures seeded in TSB at a ratio of 1:10 (gut sample: media) exhibited a growth of 3.2-fold, whereas cultures seeded in TSB supplemented with 0.5% of yeast extract (YE) at ratios of 1:5 and 1:10, experimented growths of 2.1 and 4.1-fold, respectively. We selected the latter to move forward into the fermentation process and added the cultured microbes to a new BSFL slurry. Hereafter, we will refer to the products of fermentation A and B as hydrolysate A hydrolysate B.

**Figure 1.**
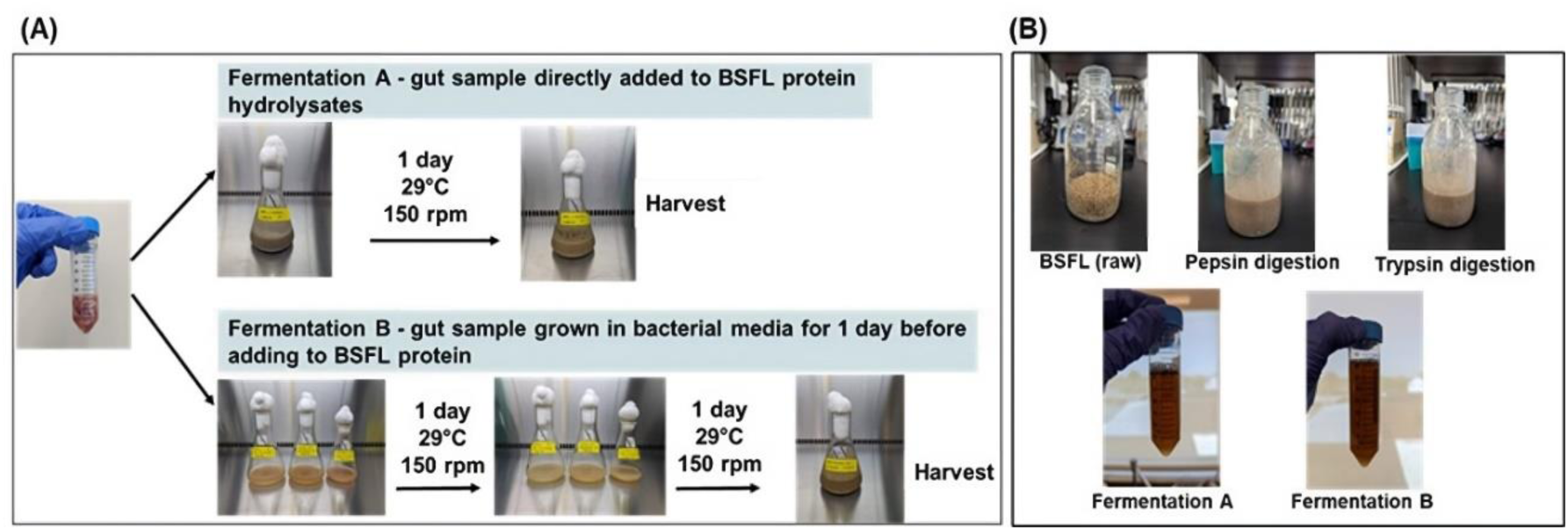
(A) Different fermentation strategies. Fermentation with microorganisms from the blue catfish gut microbiome were performed for 24 h at 29°C, and established either directly from blended gut samples (fermentation A) or after an overnight culture of gut samples in bacterial liquid media (fermentation B); (B) BSFL samples at different steps of the fermentation process.

### Isolation and composition of blue catfish gut microbiome

In order to assess the gut microbiome composition of blue catfish, we used a sample of blended gut tissue for DNA extraction and sequencing of the 16S rRNA amplicon. We obtained a total of 3135 reads belonging to 95 different OTUs (Operational Taxonomic Unit). Taxonomic identification was performed with a depth resolution of up to genus level. Table 2 is a breakdown of the bacterial genera found and their relative abundances.

**Table 2.** Gut microbiome composition of blue catfish (*Ictalurus furcatus*).^1^.

| Taxa | C | P | Percentage (%) | Taxa | C | P | Percentage (%) |
| --- | --- | --- | --- | --- | --- | --- | --- |
| Proteobacteria |  |  |  | Firmicutes |  |  |  |
| Pseudomonadaceae |  |  |  | Clostridiaceae |  | • | 8.1 |
| Pseudomonas | • | • | 6.4 | Clostridium sensu stricto 1 |  | • | 10.9 |
| Enterobacteriaceae | • | • | 4.0 | Clostridium sensu stricto 13 |  | • | 0.9 |
| Escherichia/Shigella |  |  | 3.8 | Lachnospiraceae |  |  |  |
| Plesiomonas | • |  | 2.3 | Epulopiscium |  |  | 6.4 |
| Aeromonadaceae |  |  |  | Cellulosilyticum |  |  | 2.8 |
| Aeromonas | • | • | 3.8 | Peptostreptococcaceae |  |  |  |
| Yersiniaceae |  |  | 3.5 | Romboutsia |  |  | 3.6 |
| Shewanellaceae |  |  |  | Terrisporobacter |  |  | 2.8 |
| Shewanella |  |  | 2.2 | Erysipelotrichaceae |  |  | 0.3 |
| Moraxellaceae |  |  |  | Turicibacter |  |  | 2.2 |
| Acinetobacter | • | • | 2.6 | Peptococcaceae |  |  | 0.3 |
| Xanthomonadaceae |  |  |  | Bacteroidota |  |  |  |
| Stenotrophomonas |  |  | 2.6 | Bacteroidales |  |  | 3.9 |
| Chromobacteriaceae |  |  |  | Bacteroidaceae |  |  |  |
| Crenobacter |  |  | 1.1 | Bacteroides |  |  | 3.9 |
| Chitinibacteriaceae |  |  |  | Weeksellaceae |  |  |  |
| Deefgea |  |  | 0.4 | Chryseobacterium |  |  | 0.7 |
| Sphingomonadaceae |  |  |  | Barnesiellaceae |  |  | 0.4 |
| Sphingobium |  |  | 0.3 | Sphingobacteriaceae |  |  |  |
| Fusobacteriota |  |  |  | Sphingobacterium |  |  | 0.2 |
| Fusobacteriaceae |  |  |  | Eukaryota |  |  | 2.0 |
| Cetobacterium |  |  | 6.2 | Unassigned |  |  | 1.0 |

<sup>1</sup>Percentages refer to abundance relative to the total gut microbiome at the genus level. Taxa refer to phyla (in grey), families, and genera found in gut samples, shown in no particular order. Pink and blue dots represent taxa-containing species known to produce chitinases (C) and proteinases (P), respectively (Ray *et al.*, 2012). Data was obtained from Illumina sequencing of the 16S rRNA amplicon.

The two predominant phyla detected in the blue catfish gut were Firmicutes and Proteobacteria, which accounted for 38.3% and 33.0% of the total gut microbiome, respectively. The phyla Bacteroidota and Fusobacteriota were also found with an abundance of 9.3% and 6.2%, respectively. These four phyla have been regularly reported to be primary members of fish gut microbiomes (Bledsoe et al. 2018; Ghanbari et al. 2015). Within the Firmicutes, the most abundant family was the Clostridiaceae, which alone accounted for around 20% of the total gut microbiome (Table 2), and was composed mostly of species in the *Clostridium sensu stricto* 1 cluster, of which we were able to detect *Clostridium gasigenes* (3.9% of total microbiome). *Clostridium* species are strict anaerobic fermenting bacteria commonly found as commensals in both land animal and fish guts (Bledsoe et al. 2018; Egerton et al. 2018; Liu et al. 2016; Wang et al. 2018). Within the Proteobacteria, the most abundant genera detected were *Pseudomonas* (6.4%), *Aeromonas* (3.8%) and members of the Enterobacteriaceae such as *Escherichia/Shigella* (3.8%) and *Plesiomonas* (2.3%), all of whom have been previously reported to be abundant in the intestinal microbiota of freshwater fish species (Sugita et al. 1995; Wang et al. 2018).

Chitinases and proteases are enzymes important for the digestion of small invertebrates and insects such as BSFL. Chitin is the second most abundant polysaccharide in nature and the main constituent in the exoskeleton of insects and water invertebrates. Our study found blue catfish contains strains belonging *to Enterobacter, Aeromonas, Pseudomonas, Plesiomonas* and *Acinetobacter* (Table 2), which all represent genus with species commonly reported to degrade chitin and to be present in the gut of carnivore, omnivore and zoo-planktivore fish (Egerton et al. 2018; Ray et al. 2012). *Cetobacterium* was also detected by our study to be abundant in blue catfish (6.2% of total microbiome) and has also been reported to be dominant in carnivorous and omnivorous fish microbiomes when compared to herbivorous fish (Liu et al. 2016). Species of the *Enterobacter, Pseudomonas, Aeromonas* and *Acinetobacter* have also been found to produce proteases which aid the breakdown of proteins into amino acids in the fish intestine (Ray et al. 2012). While *Clostridium* is mostly associated with cellulase and amylase activity in herbivore species, it has also been reported to have the ability to break down proteins (Liu et al. 2016; Ray et al. 2012).

### Changes in microbiome composition through cultivation and fermentation

Extraction and cultivation of the gut microbiome are bound to alter the composition of the existing bacterial community, as the culture conditions imposed will favor the growth and survival of some species over others. In fact, most intestinal bacteria are widely considered to be unculturable and have never been isolated in the laboratory (Stewart 2012). In order to monitor these changes, we assessed the microbial composition of the BSFL slurry samples at different stages of the fermentation process by DNA extraction and sequencing of the 16S rRNA amplicon. Samples were analyzed before the addition of gut microbes and after the 24 h fermentation steps A and B. We also analyzed the effect of culturing gut microbes in the liquid bacterial medium for 24 h (TSB supplemented with 0.5% of YE at a 1:10 seeding ratio) prior to fermentation B (Table 3).

**Table 3.** Alpha diversity of the bacterial communities at different points of the BSFL fermentation process.^1^.

| Sample | Reads | Alpha diversity |  |  |  |  |
| --- | --- | --- | --- | --- | --- | --- |
|  |  | Observed | ace | Chao1 | Simpson | Shannon entropy |
| BSFL slurry | 12272 | 310 | 310 | 310 | 0.976 | 0.645 |
| Blue catfish gut | 3135 | 95 | 95 | 95 | 0.977 | 0.579 |
| Fermentation A | 1514 | 44 | 44 | 44 | 0.936 | 0.454 |
| Gut bacterial culture | 1738 | 49 | 49 | 49 | 0.948 | 0.470 |
| Fermentation B | 2007 | 60 | 60 | 60 | 0.959 | 0.505 |

<sup>1</sup>Different parameters were used for the assessment of species richness and evenness of the samples: observed, ace, chao1, simpson and shannon indexes.

We observed a higher number of overall reads in the BSFL slurry sample before fermentation, than in the samples taken from blue catfish gut and during the fermentation processes (Table 3). Surprisingly, alpha diversity parameters that measure species richness such as observed richness, ace and chao1, were also higher for BSFL slurry before fermentation than for the rest of the samples. Alpha diversity parameters that consider both species richness and evenness, such as the simpson and shannon indexes, were more even, but we also observed a higher shannon value for the initial BSFL slurry (Table 3). As expected, the bacterial composition of the BSFL slurry before fermentation was very different from that in the blue catfish microbiome (Figure 2). The microbial taxa identified in the BSFL slurry are likely a representation of the bacterial community present in the dry BSFL material but could also come from other non-related sources since enzymatic digestions were not conducted in sterile conditions. Previous reports have shown that the most dominant bacterial genera in BSFL gut microbiota include *Morganella*, *Providencia*, *Dysgonomonas*, *Ignatzschineria*, *Enterobacter*, *Proteus*, *Enterococcus*, *Bacillus*, *Klebsiella*, *Citrobacter*, *Scrofimicrobium* and *Actinomyces* (Eke et al. 2023). We found our BSFL slurry to be specially rich in Cyanobacteria (47%), but likewise we were able to find *Bacillus* (6%), *Dysgonomonas* (3.6%), *Enterococcus* (2.7%), *Providencia* (1.3%), *Actinomyces* (0.5%), *Morganella* (0.2%), *Proteus* (0.2%), and other previously non reported genera such as *Caldalkalibacillus* (2.5%), *Gilliamella* (1.2%), *Lachnoclostridium* (1%) and *Terrisporobacter* (1%). Most of these genera, with the exception of Enterococcus, were reduced significantly during the fermentation process.

**Figure 2.**
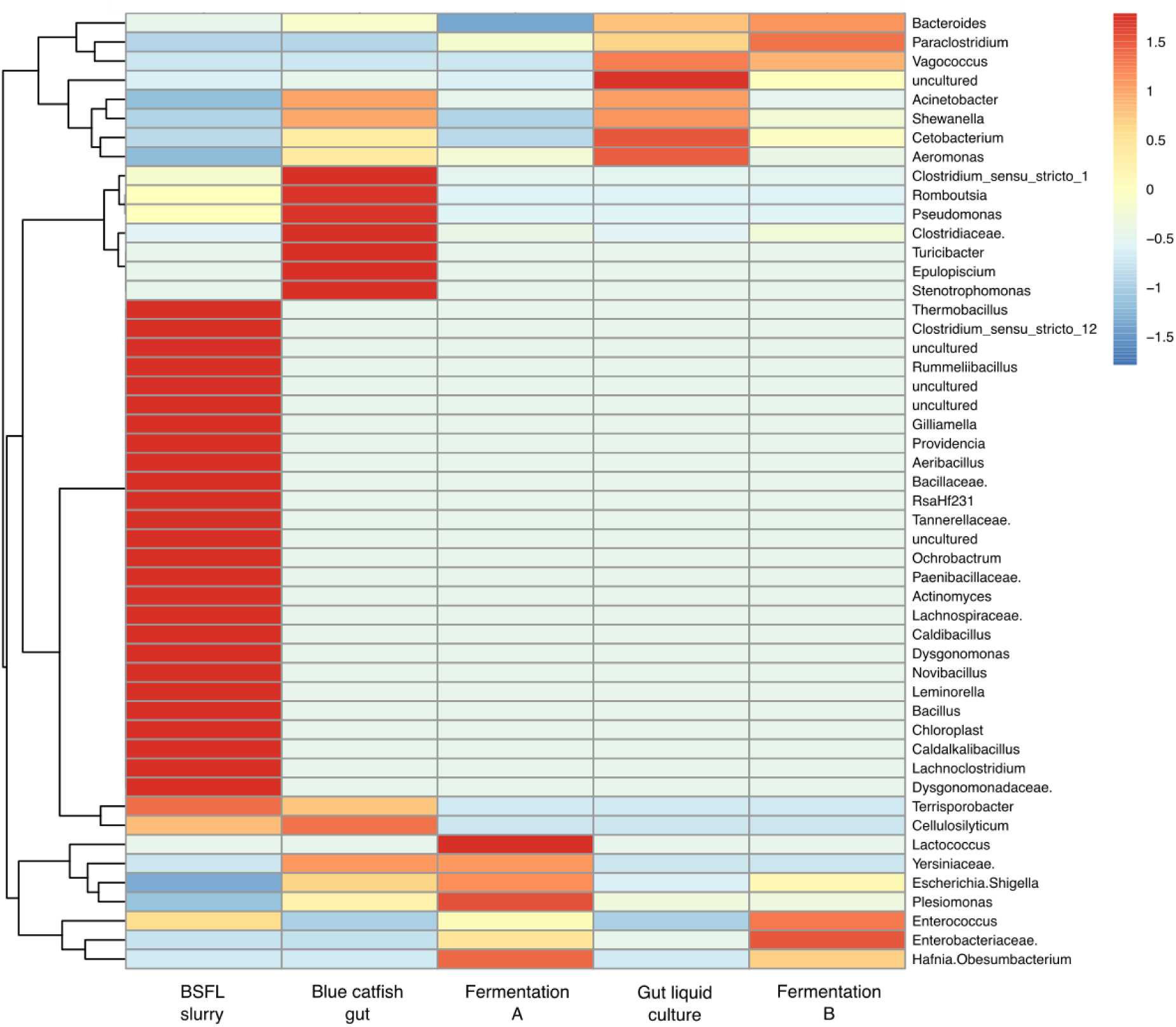
Bacterial genus abundance heatmap at different points of the BSFL fermentation process. The top 50 identified taxa are shown. Color represents the abundance of each taxon, and vertical clustering indicates the similarity of the abundance between different taxa. The absolute value of the legend represents the distance between the raw score and the mean population of the standard deviation. The legend is negative when the raw score is below the mean.

As anticipated, many of the bacterial genera identified in the blue catfish gut were lost through cultivation and fermentation (Figure 2). Species richness decreased by approximately 2-fold in fermentation A and by 1.6-fold in fermentation B (Table 3). Lost bacterial genera included important chitinase and protease producing genera such as *Pseudomonas* and members of the Clostridiaceae (including *Clostridium*), and extended to the *Chryseobacterium, Sphingobacterium, Turicibacter, Cellulosyticum, Epulopiscium, Romboutsia, Terrisporobacter, Deefgea, Crenobacter, Stenotrophomonas* and members of the Erysipelotrichaceae and Peptococcaceae families. Conversely, we observed an increase of *Paraclostridium* and members of the Enterobacteriaceae in both fermentation strategies. In comparison, fermentation A favored the growth of *Escherichia/Shigella, Plesiomonas* and Yersiniaceae, whereas fermentation B favored the growth of Bacteroides and other Bacteroidales, *Cetobacterium, Shewanella* and members of the Barnesiellaceae. Cultivation of gut samples in TSB supplemented with YE was able to sustain the growth of *Cetobacterium, Aeromonas, Shewanella, Escherichia/Shigella, Plesiomonas, Acinetobacter, Vagococcus, Paraclostridium, Bacteroides* and other Bacteroidales, *Enterobacter* and other members of the Enterobacteriaceae, and members of the Barnesiellaceae.

### Effect of fermented hydrolysates in in-vitro fish cell proliferation

In order to assess if products of the fermentation of BSFL hydrolysates with catfish gut microbiota can be used to replace fetal bovine serum (FBS) in *in vitro* fish cell cultivation, we tested their effect on the proliferation of ZEM2S zebrafish (*D. rerio*) embryo fibroblasts. Previous reports show that low concentrations of BSFL protein hydrolysates (0.001 - 0.1 mg/mL) are able to support ZEM2S cell growth in media containing low concentrations of FBS (1 - 2.5%) (Batish et al. 2022). Based on these results, we chose to test a range of our hydrolysate concentrations (0.001, 0.01, 0.1, 1 mg/mL) and FBS concentrations (0, 1, 2.5, 10%) for their effect on cell proliferation and morphology over three days. Our hydrolysate concentrations were based on the peptide concentrations obtained by the *o*-phthalaldehyde (OPA) methodology.

As expected, decreasing the concentration of FBS in the media resulted in a linear decrease in cell density and had a detrimental impact on cell morphology. Compared to cell cultures grown in basal medium containing 10% FBS, cultures grown with 2.5%, 1% and 0% FBS showed cell proliferation reductions of 28.1%, 53.9% and 75.8% after 72 hours, respectively (Figure 3A, S1). Also, as previously reported, cells lost their original spindle-like fibroblast morphology at low FBS concentrations and started exhibiting starved characteristics as they became thinner, spikier, more elongated, and three-dimensional (Figure S1) (Batish et al. 2022). The severity of these cell morphology alterations was proportional to the decrease in FBS concentration.

**Figure 3.**
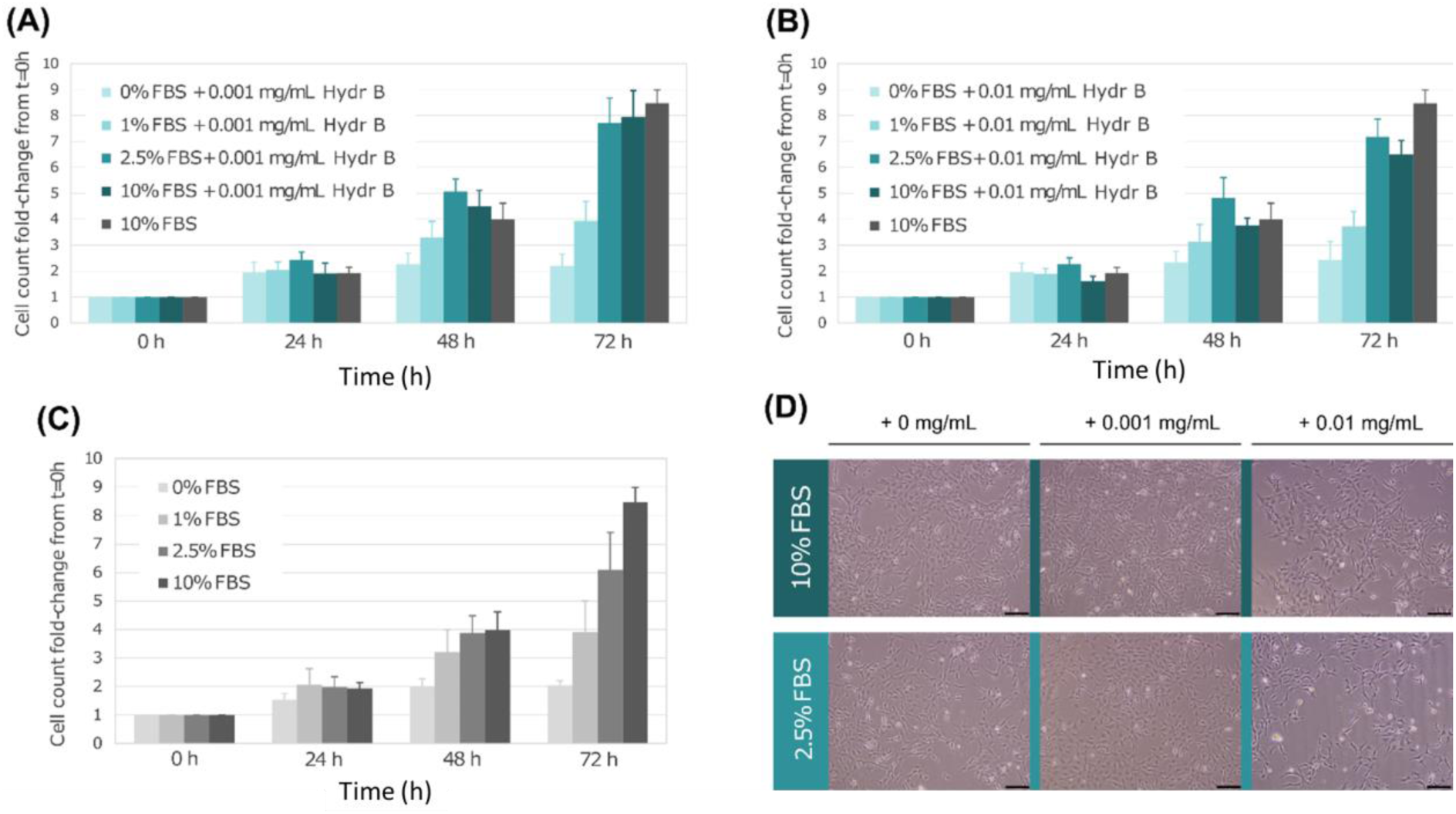
Effect of supplementation of hydrolysate B on ZEM2S *in vitro* proliferation at decreasing FBS concentrations. Cell proliferation in media supplemented with (A) 0.001 mg/mL, (B) 0.01 mg/mL; (C) no hydrolysate B over three days; (D): Cell morphology at different conditions. Different hues of blue represent different FBS concentrations (0%, 1%, 2.5%, 10%). All values are fold-change increases calculated over the first time point (0 h), hence all values at 0 h start as 1. Bars represent the average and error bars represent the standard deviation between 4 biological replicates.

We then proceeded to test the performance of our fermented hydrolysates, starting with hydrolysate B. At high peptide concentrations (0.1 and 1 mg/mL), supplementation of hydrolysate B resulted in cytotoxic effects on the cells after 72 hours of culture (Figure S2, S3). These effects negatively impacted cell number and morphology, increased with hydrolysate concentration, and were observed in all FBS concentrations assayed including the standard 10% FBS concentration (Figure S2, S3). At lower peptide concentrations (0.01 and 0.001 mg/mL), supplementation of hydrolysate B only seemed to improve cell proliferation in cultures with 2.5% FBS to levels comparable with the 10% FBS control (Figure 3). However, the difference between 2.5% FBS cultures with or without hydrolysate B after 72 hours did not appear to be statistically significant, even upon experimental repeat. Similar proliferation patterns were obtained when supplementing with hydrolysate A. Here, supplementation of 0.01 mg/mL in cultures at 2.5% FBS resulted in an apparent similar cell proliferation improvement (Figure 4), but likewise, this difference was not statistically supported. In any case, similar results were obtained when supplementing the cultures with BSFL hydrolysate controls taken previous to bacterial fermentation (Figure 4C), suggesting fermentation would not be responsible for the proliferation improvements observed. Additionally, we did not observe any noticeable changes in cell morphology between cultures with and without hydrolysates A and B at concentration ranges of 0.01 and 0.001 mg/mL.

**Figure 4.**
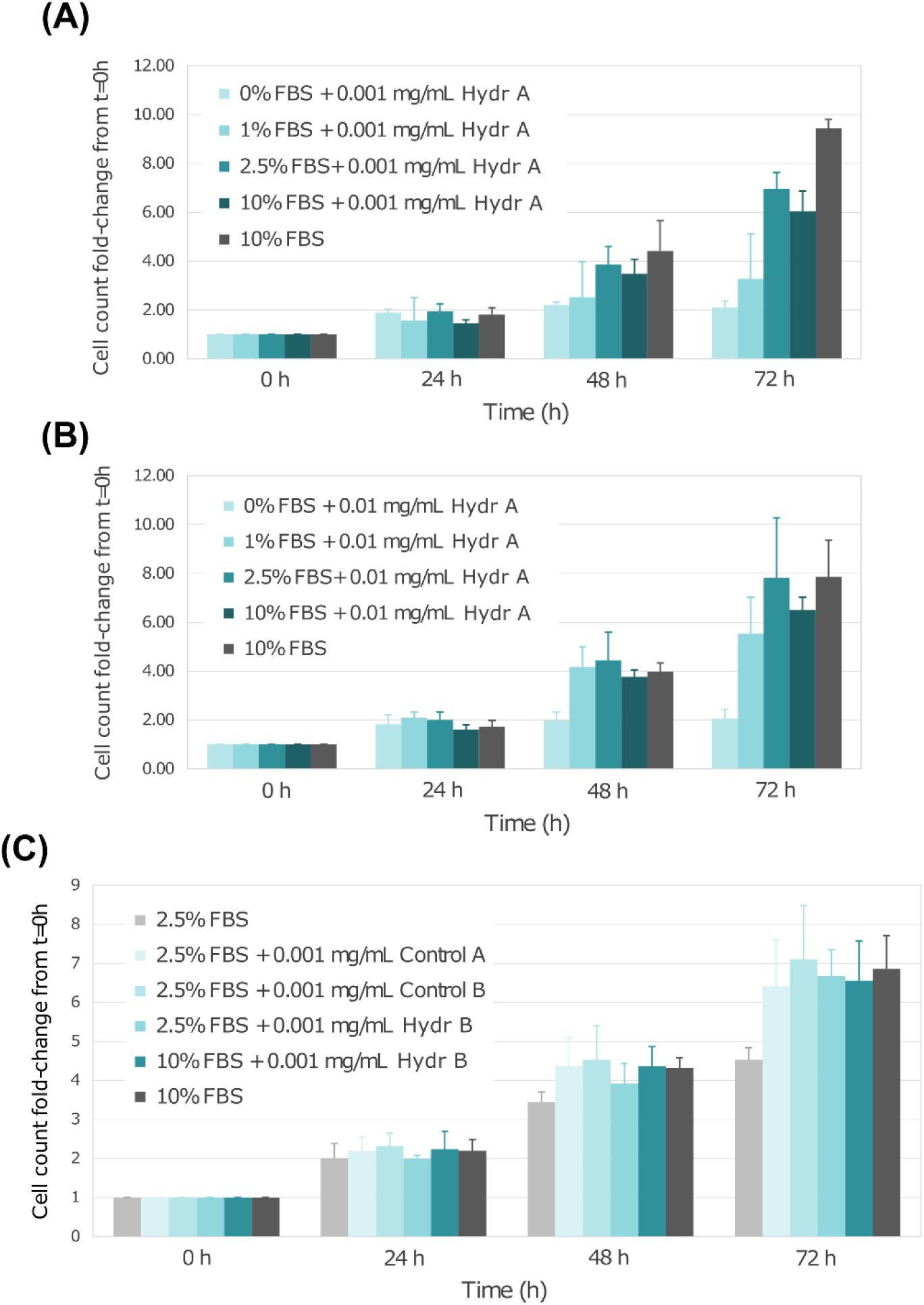
Effect of supplementation of hydrolysate A on ZEM2S in vitro proliferation at decreasing FBS concentrations. Cell proliferation in media supplemented with (A) 0.001 mg/mL and (B) 0.01 mg/mL of hydrolysate A over three days. Different hues of blue represent different FBS concentrations (0%, 1%, 2.5%, 10 %). (C) Cell proliferation in media supplemented with BSFL hydrolysate controls taken previous to fermentation with blue catfish gut microbes. All values are fold-change increases calculated over the first time point (0 h); hence, all values at 0 h start as 1. Bars represent the average and error bars represent the standard deviation between 4 biological replicates.

## Conclusion

Substituting FBS-containing media with serum-free media is a key focus in cellular agriculture research. Numerous researchers have explored various alternatives, such as free amino acids, protein hydrolysates, recombinant proteins, and intact protein isolates, to replace serum in cell culture media. While certain proteins and peptides have proven successful in replacing or reducing serum in cell culture media, feedstock bioconversion through gut microbial communities has not been explored in the cultivated meat industry. In this study, we implemented a nature-inspired bioconversion approach, leveraging the gut microbial communities of insectivorous fish (Blue catfish), to convert black soldier fly larvae, one of the most sustainable and abundant insects, into postbiotics for the development of cell culture media. Fermentation with gut microbial communities significantly increased the degree of hydrolysis up to 16.1%. Our culture method favored the growth of important protease and chitinase-producing genera like *Aeromonas, Acinetobacter, Plesiomonas*, and members of the Enterobacteriaceae family but lost important genera like *Pseudomonas* and *Clostridium*. Briefly, microbiome composition alterations were observed during fermentation of BSFL protein hydrolysates: *Pseudomonas* was lost, *Clostridium* was heavily reduced, *Aeromonas* and *Acinetobacter* were slightly reduced but maintained, and *Paraclostridium* and members of the Enterobacteriaceae increased. Fermentation also favored the growth of other genera like *Plesiomonas* and *Cetobacterium*.

Following the incorporation of postbiotics into the ZEM2S cell culture media, there was a notable enhancement in cell proliferation observed at 2.5% serum concentrations, with comparable proliferation levels observed at 10% serum. In contrast to our prior investigations involving the application of protein hydrolysates from BSFL, the improvement in cell proliferation with postbiotics was higher, while it was not statistically significant. Further investigations are essential to assess the postbiotic composition, including short-chain fatty acids, glucose, vitamins, minerals, etc., and their influence on cell metabolism through metabolomics.

## Acknowledgments

This research was financially supported by the Agriculture and Food Research Initiative (AFRI) Sustainable Agricultural Systems program, grant no. 2021-699012-35978 from the USDA National Institute of Food and Agriculture, and Texas A&M AgriLife Research. The authors would like to gratefully acknowledge Mohammad Zarei for his support.

## Notes

### Competing Interest Statement

The authors have declared no competing interest.

