## Supplemental Data for "Black Soldier Fly Bioconversion to Cultivated Meat Media Components Using Blue Catfish Gut Microbiome"

**Figure S1. Effect of FBS reduction on ZEM2S *in vitro* cell proliferation. (A)** Cell number counts over time (0, 24, 48, 72 h) in ZEM2S cultures with decreasing FBS concentrations (0%, 1%, 2.5%, 10%). All values are fold-change increases calculated over the first time point (0 h), hence all values at 0 h start as 1. Bars represent the average and error bars represent the standard deviation between 4 biological replicates. **(B)** Phase contrast microscopy of ZEM2S cultures after 72 h in different FBS concentrations. Cell morphology and density is affected as FBS concentrations decrease. Scale bar, 100 µm.

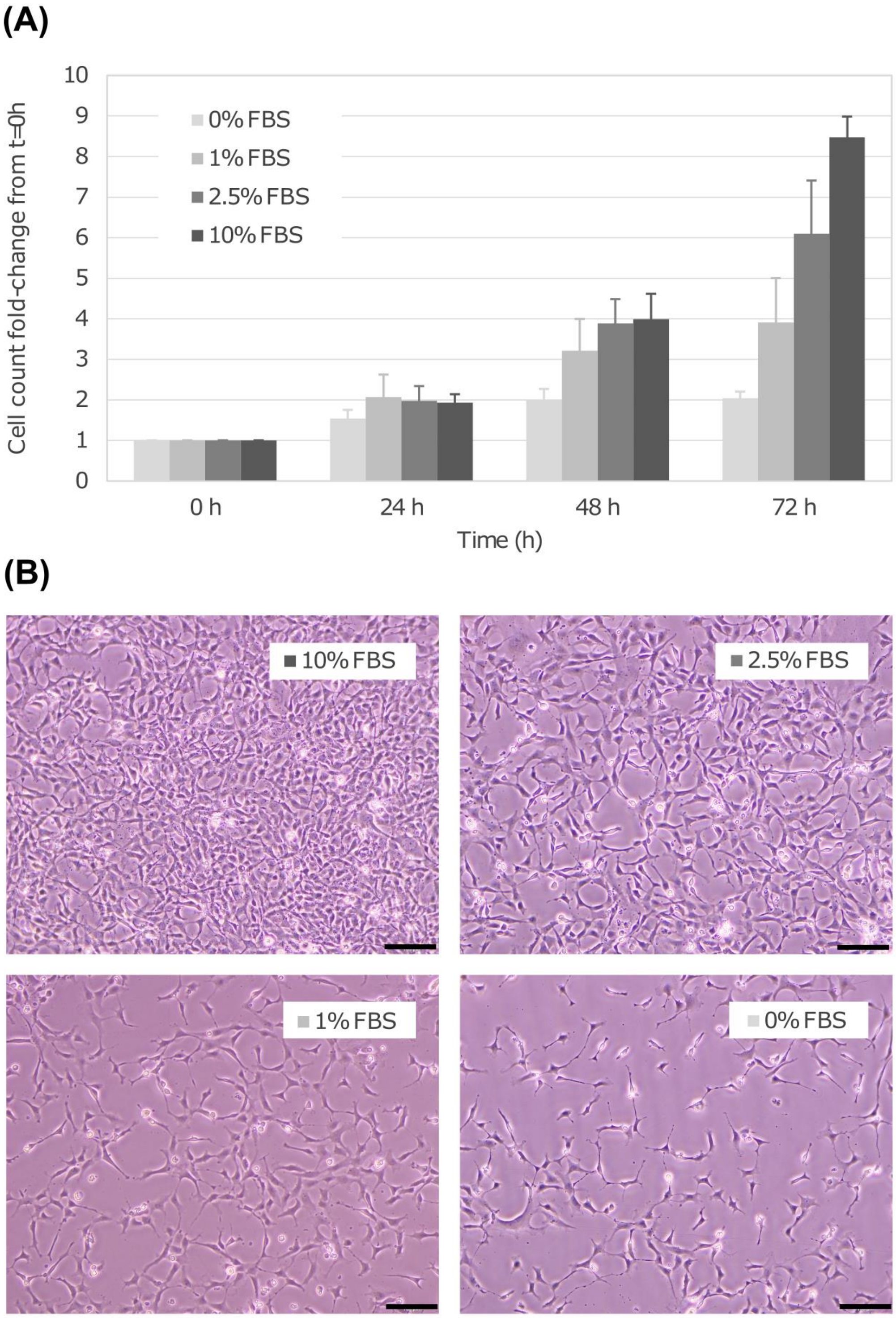

**Figure S2. Effect of supplementation of high concentrations of hydrolysate B on ZEM2S *in vitro* proliferation at decreasing FBS concentrations.** Cell proliferation in media supplemented with **(B)** 0.1 mg/mL, **(C)** 1 mg/mL and **(A)** no hydrolysate B over three days. Different hues of blue represent different FBS concentrations (0%, 1%, 2.5%, 10%). All values are fold-change increases calculated over the first time point (0 h), hence all values at 0 h start as 1. Bars represent the average and error bars represent the standard deviation between 4 biological replicates.

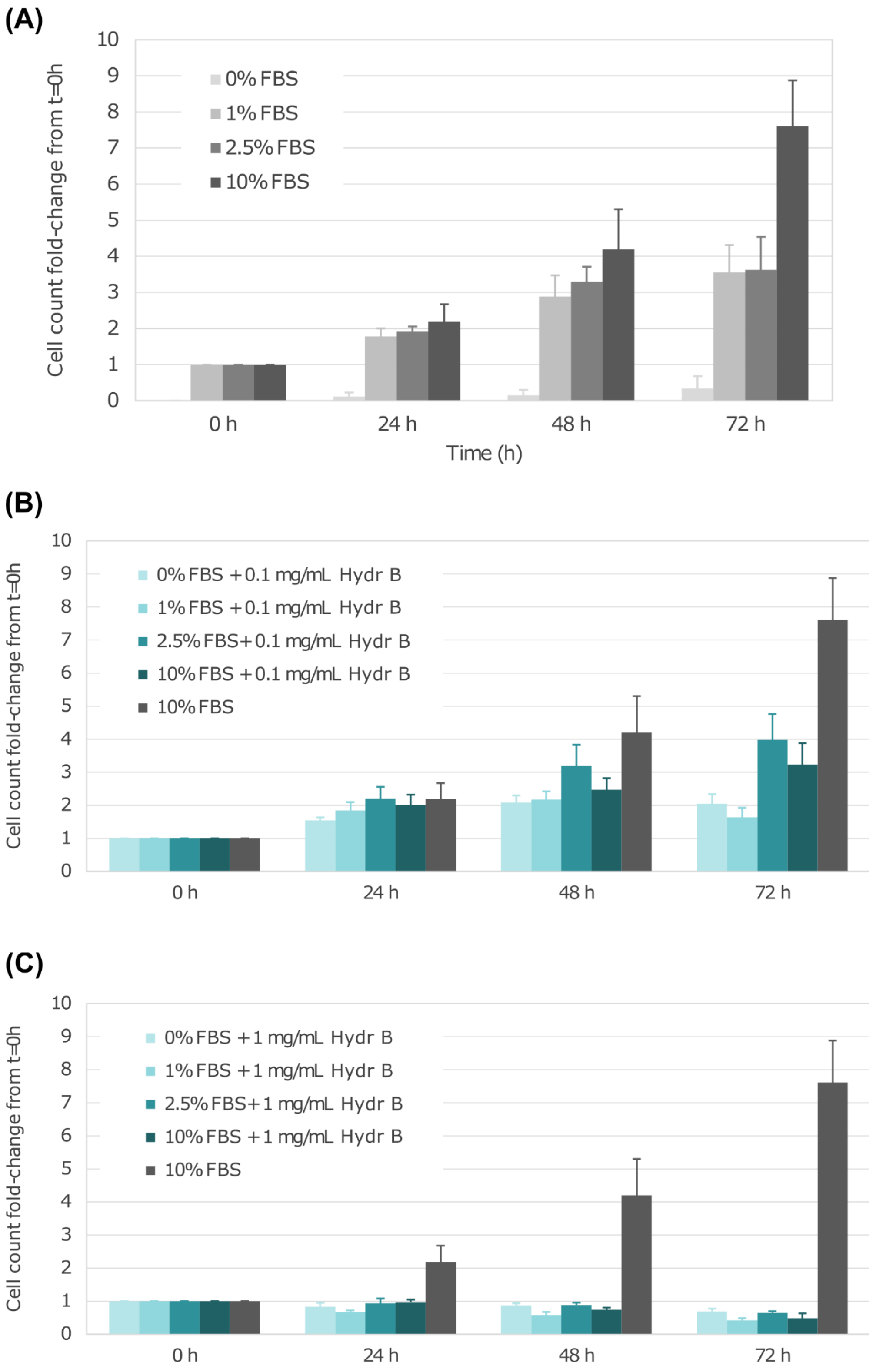

**Figure S3. Cell morphology after supplementation with high concentrations of hydrolysate B at decreasing FBS concentrations.** Phase contrast microscopy of ZEM2S cultures after 72 h in media supplemented with 0 mg/mL, 0.1 mg/mL and 1 mg/mL of hydrolysate B. Different hues of gray represent decreasing FBS concentrations. Cell morphology and density is affected as FBS concentrations decrease and hydrolysate B concentrations increase. Scale bar, 100 μm.

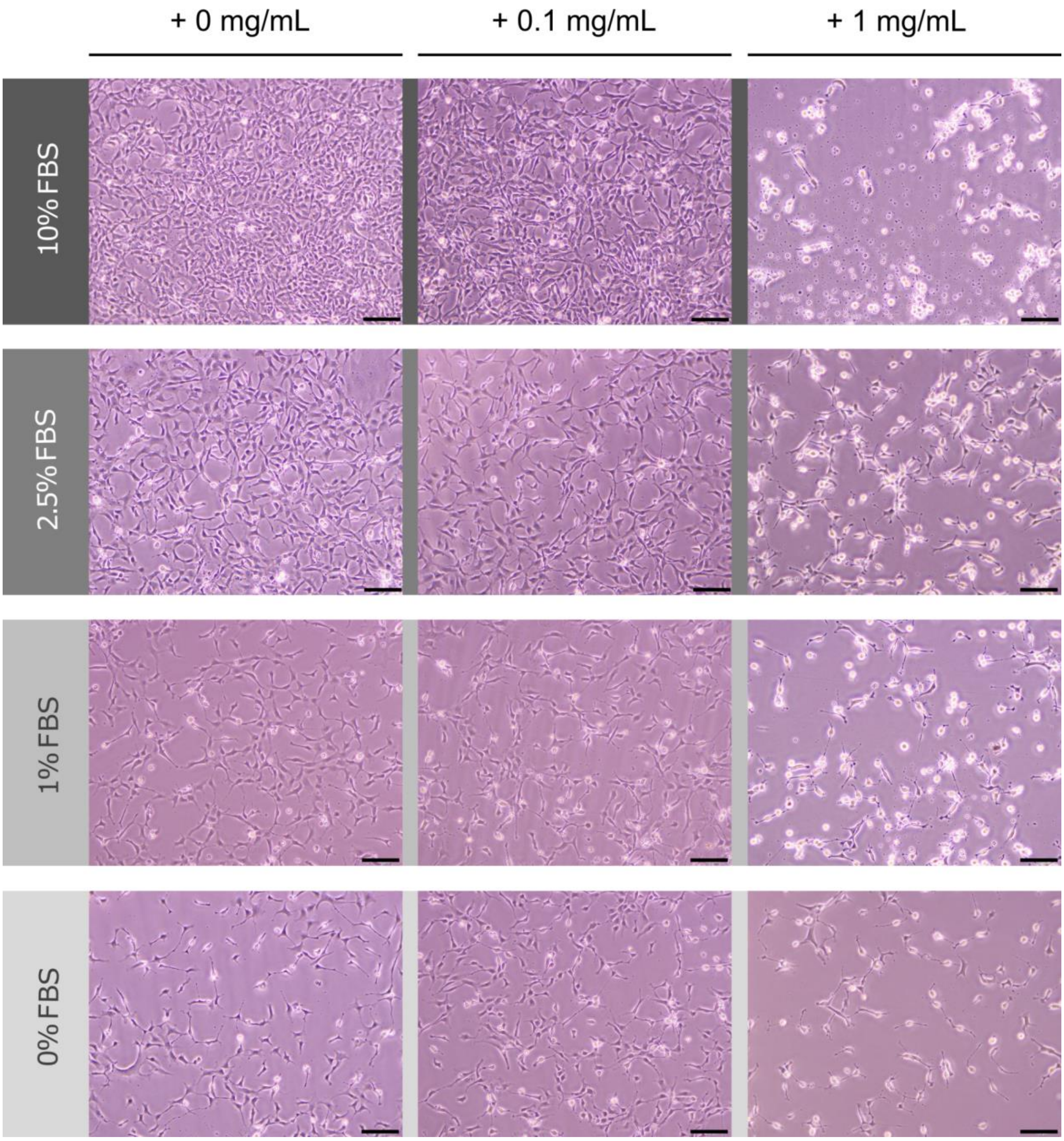
